# The Flowering Repressor SVP recruits the TOPLESS co-repressor to control flowering in chrysanthemum and *Arabidopsis*

**DOI:** 10.1101/2021.11.23.469726

**Authors:** Zixin Zhang, Qian Hu, Yuqing Zhu, Zheng Gao, Erlei Shang, Gaofeng Liu, Weixin Liu, Rongqian Hu, Xinran Chong, Zhiyong Guan, Weimin Fang, Sumei Chen, Bo Sun, Yuehui He, Jiafu Jiang, Fadi Chen

**Affiliations:** State Key Laboratory of Crop Genetics and Germplasm Enhancement, the Key laboratory of Landscaping, Ministry of Agriculture, College of Horticulture, Nanjing Agricultural University, Nanjing 210095, China; Shanghai Center for Plant Stress Biology, Chinese Academy of Sciences Center for Excellence in Molecular Plant Sciences, Shanghai 201602, China; State Key Laboratory of Pharmaceutical Biotechnology, School of Life Sciences, Nanjing University, Nanjing 210023, China; Peking University Institute of Advanced Agricultural Sciences, Weifang, Shandong 261000, China

**Keywords:** Chrysanthemum, *Arabidopsis*, flowering time, protein interaction, co-repressor, *FLOWERING LOCUS T*.

## Abstract

Plant flowering time is a consequence of the perception of environmental and endogenous signals. The MCM1-AGAMOUSDEFICIENS-SRF-box (MADS-box) gene SHORT VEGETATIVE PHASE (SVP) is a pivotal repressor that negatively regulates the floral transition during the vegetative phase. The transcriptional corepressor TOPLESS (TPL) plays critical roles in many aspects of plant life. An interaction first identified between the second LXLXLX motif (LRLGLP) of CmSVP with CmTPL1-2, which can repress the expression of a key flowering factor *CmFTL3* by binding its promotor CArG element in chrysanthemum. Genetic analysis suggested that the CmSVP-CmTPL1-2 transcriptional complex is a prerequisite for SVP to act as a floral repressor, which reduces *CmFTL3* transcriptional activity. *CmSVP* rescued the phenotype of the *svp*-*31* mutant in *Arabidopsis*, and overexpression of *AtSVP* or *CmSVP* in the *Arabidopsis* dominant negative mutation *tpl*-*1* led to a loss-of-function in late flowering, which confirmed the highly conserved function of *SVP* in the two completely different species. Thus, we have validated a conserved machinery wherein SVP relies on TPL to inhibit flowering through the direct regulation of *FT*, which is more meaningful for the evolution of species and could be translated to high-quality cultivation and breeding of crops.

## Introduction

In plants, the regulation of multiple endogenous cues as well as the response to the external environment is the crucial criteria for activation of flowering time (Kinoshita and Richter, 2020). Serveral flowering-regulated MADS-box genes have been studied in recent years, and yet poorly negative regulators are identified and understood in plants. SHORT VEGETATIVE PHASE (SVP), as one transcription factor of the MCM1-AGAMOUSDEFICIENS-SRF (MADS)-box gene family, can respond to themosensory, gibberellin, and autonomous pathways (Andrés et al., 2014; Fernández et al., 2016). An analyses of evolution showed that the SVP are highly conserved and present in nearly all eudicot species (Liu et al., 2018).

Mutants of *SVP* show a phenotype of early flowering under long days (LDs) or short days (SDs) that represses the expression of *FLOWERING LOCUS T* (*FT*), *TWIN SISTER OF FT* (*TSF*), and *SUPPRESSOR OF OVEREXPRESSION OF CONSTANS1* (*SOC1*) to maintain the vegetative phase of plants (Andrés et al., 2014; Hartmann et al., 2000; Jang et al., 2009; Li et al., 2008). The study of *Arabidopsis* indicated that SVP could bind to the CC (A/T) _6_ GG (CArG) in the promoter region and form a dimer complex to play a regulatory function (Folter and Angenent, 2006; Gregis et al., 2013; Hartmann et al., 2010), but the negative action controlled by this MADS-domain transcription factor is unclear.

And yet a transcriptional co-repressor TOPLESS (TPL)/TPL-RELATED (TPR) is also involved in a set of proteins responsible the switch from vegetative to reproductive phase by inhibiting transcription of *FT* (Causier et al., 2012; Goralogia et al., 2017; Krogan et al., 2012; Zhang et al., 2019). TPL/TPR family proteins, as universal transcription GRO/Tup1-like co-repressors, are widely present in plants and participate in the biological processes of growth and development, such as plant hormone signaling pathways, various stress responses, and cycle rhythm clock regulation (Causier et al., 2012; Plant et al., 2021). TPL/TPR protein is able to interact with specific transcription factors directly or indirectly, thereby inhibiting the expression of target genes and the performance of the signal transduction pathway (Pauwels et al., 2010). The mutation (*tpl*-*1*) at the 176^th^ amino acid position of the N-terminal domain of the TPL protein from aspartic acid to histidine is a dominant negative mutation that can cause dramatic temperature-sensitive abnormalities in growth and development, and TPL has been confirmed as a transcriptional co-repressor that interacts with the EAR domain of IAA12/BDL to control *ARF* transcriptional activity in an auxin-dependent manner (Long et al., 2006). Studies have suggested that TPL/TPR genes have a function during the transition to flowering(Leydon et al., 2021). Barry (Causier et al., 2012) found that the delayed flowering caused by TOE1’s inhibition of *FT* expression is dependent on TPL/TPR. TPL can interact with CO through the microprotein miP1a/b, thereby inhibiting the expression of *FT* and delaying the flowering of *Arabidopsis* (Graeff et al., 2016). Further research has revealed the function of the microprotein miP1a in floral repression, in which a repressor complex with miP1a/b, CO/CO-like transcription factors, TPL, and JMJ14 prevents flowering by repressed *FT* gene transcription in *Arabidopsis* (Rodrigues et al., 2021). The dominant negative *tpl*-*1* mutant sequence is driven by the endogenous gene *SUC2* promoter, which results in the flowering time being significantly earlier in *Arabidopsis*. The study further speculated that CYCLING DOF FACTOR 1 is combined with TPL to regulate the expression of *CO* and *FT*, thus, inhibiting flowering in the photoperiod pathway (Goralogia et al., 2017). A recent study found that GA signals could be mediated by the GAF1-TPR complex to repress the expression of *ELF3*, *SVP*, and TEMs, which leads to the induction of *FT* and *SOC1* (Fukazawa et al., 2021).

Chrysanthemum is one of the most important ornamental plants used worldwide, and it is widely cultivated as cut, potted, and garden flowers, and the flowers of some cultivars are a resource for medicinal materials (Teixeira, 2003). Therefore, the ornamental and commercial values are dependent on the appropriate time of year. And the discovery of key genes involved in the vegetative stage and responses to temperature or light have great significance for generating new chrysanthemum cultivars and ensuring year-round production. In previous studies, *Arabidopsis FT* homologous genes *CsFTL1*, *CsFTL2*, and *CsFTL3* were cloned from *Chrysanthemum seticuspe* and *CsFTL3* was further elucidated as a key factor in the photoperiod pathway of chrysanthemums (Oda et al., 2012). The CONSTANS homologous gene *CmBBX8* belonging to the BBX family isolated from a day-neutral chrysanthemum ‘Yuuka’ accelerates flowering by targeting *CmFTL1* directly (Wang et al., 2020). Another member of the BBX family in chrysanthemums, *CmBBX24* suppresses flowering time by inhibiting GA biosynthesis (Yang et al., 2014). The *TERMINAL FLOWER 1* homologous gene *CsAFT* can inhibit flowering through the disruption of the FT-FD complex (Higuchi et al., 2013). *CmNF-YB8* is involved in the age pathway to accelerate the transition from the juvenile to adult phase in chrysanthemums (Wei, Y. et al., 2017). We previously identified a highly homologous gene to *SVP* in the chrysanthemum ‘QD026’, which we named *SVP1* (CL11972. Contig2_All), and its expression level declined while that of *FT* increased based on a result of RNA-seq during floral transition (Cheng et al., 2018). In the present study, *CmSVP* was cloned from the chrysanthemum ‘Jinba’ and the transgenic chrysanthemum was generated. The expression of *CmFTL3* was detected, respectively, in overexpression and knockdown lines of *CmSVP*, which suggests that the expression pattern of *CmFTL3* is negatively regulated by *CmSVP*. Evidence from the genetic and ChIP assay showed that CmSVP is responsible for the reduction in *CmFTL3* transcription directly. Then, we showed that CmSVP recruits CmTPL1-2 to reduce *CmFTL3* transcription in the chrysanthemum ‘Jinba’, and the mechanism is also conservative in *Arabidopsis*. The combined data reveal that SVP repressing flowering to keep the plants in a vegetative stage depends on TPL activity. As *Arabidopsis* is a facultative long-day plant, and chrysanthemum ‘Jinba’ is a strict short-day variety that takes on a complex hexaploid, a highly conserved SVP-TPL machinery is significance for the species evolution. And moreover, These findings will help to elucidate the functions of numerous orthologous and homologous genes in floral transition in plants.

## Results

### *CmSVP* delays the transition to flowering in chrysanthemum

To determine the function of *CmSVP* in the chrysanthemum ‘Jinba’, the length of *CmSVP*, which was cloned from ‘Jinba’, consists of a 669 bp coding sequence and encodes 223 amino acids with a predicted molecular mass of 25.46 kDa and a *pI* (isoelectric point) of 6.797. The sequence features and expression pattern of CmSVP in chrysanthemums were initially verified (Fig. S1). The sequence features a conserved MADS domains at the N termini, which contain the non-translatable binding site of miR396 (Fig. S1A). As the *SVP* mRNA in *Arabidopsis* decays, it is triggered by miR396(Palatnik et al., 2003; Yang et al., 2015). CmSVP-mut396 with four mismatches in the core pairwise region of miR396 was obtained by site-directed mutagenesis to avoid the miR396-mediated translation inhibition (Fig. S1A). As tested by the yeast hybrid system, CmSVP is not related to any of the transcriptional autoactivation activities, while the VP16 is a fragment of viral DNA sequence that can reverse the feature (Fig. S1B). Evolutionary analysis showed that CmSVP is closely related to AaSVP, which is derived from *Artemisia annua* (Fig. S1C, Fig. S2). Moreover, the specific sequence information is shown in Fig. S2. The relatively high abundance of the transcript was present in stems of the vegetative phase, followed by the leaves and buds at the reproductive stage (Fig. S1D). In addition, laser confocal microscopy was employed to reveal the GFP signals of CmSVP-GFP fusion gathered in the nuclei, while those of control 35S::GFP were presented in the whole cell (Fig. S1E).

In the chrysanthemum ‘Jinba’, a strategy that involves the recruitment of additional VP64 for 35S::*CmSVP*-*VP64* fusion was used to reverse its transcriptional repressor activity. VP64 is the four tandem repeats of VP16, which is capable of turning a repressive transcription regulator into an activator completely. And may causes a similar or stronger phenotype compared to the knockout plants(Guo et al., 2018; Suzuki et al., 2014; Triezenberg et al., 1988). Three transgenic lines (H-2, E-3, and I-5) of 35S::*CmSVP*-*VP64* were selected that flowered significantly earlier than the wild type (WT) plant ‘Jinba’, while 35S::*CmSVP* (OX) (A-1, B-2, and Z-2) showed a later flowering time compared to WT ‘Jinba’ (Fig. 1A, B, Fig. S3) with molecular identification (Fig. S4A, Fig. 1C). The first involucral primordia was initiated in 35S::*CmSVP*-*VP64* plants, 14 days after being transplanted and grown under SD conditions. This developmental stage was 13 days earlier than in WT plants. When the 35S::*CmSVP*-*VP64* plants had already reached the open-flower stage, the OX plants were still at the bud formation stage, which was 20–23 days later than that in WT plants (Fig. 1B). We next investigated whether the expression of *CmFTL3* changed in transgenic plants. Compared to the WT ‘Jinba’, 35S::*CmSVP* plants showed dramatically decreased levels of *CmFTL3* mRNA, while 35S::*CmSVP*-*VP64* plants presented an increasing trend (Fig. 1C).

**Fig. 1.**
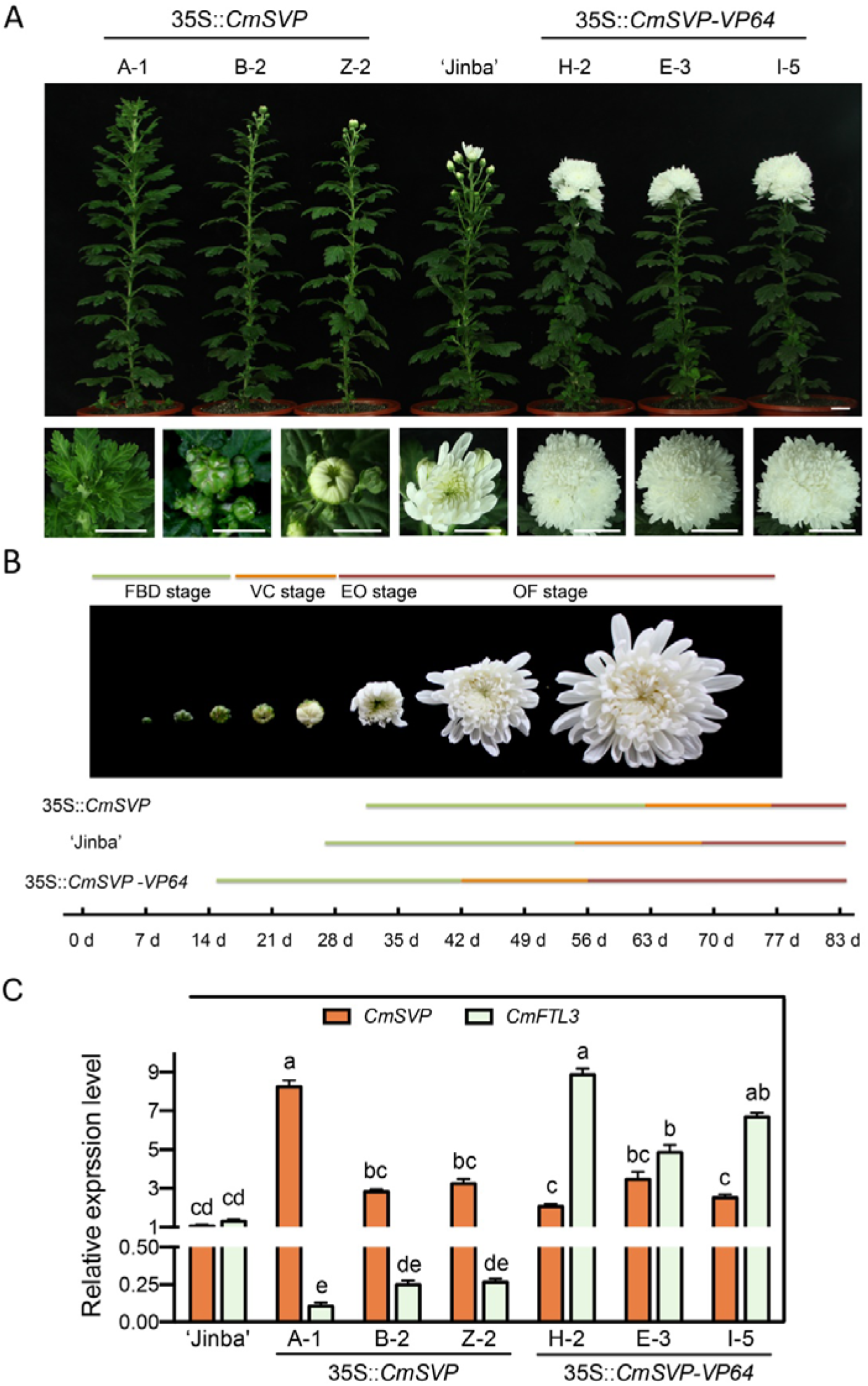
Phenotypes of *CmSVP* transgenic ‘Jinba’ plants. A. The phenotypic consequence of overexpression (A-1, B-2, and Z-2) and a constitutively active form of *CmSVP* (H-2, E-3, and I-5). Scale bar = 1.5 cm. B. Developmental process of flower buds in wild type ‘Jinba’ and transgenic plants. Flowering time was calculated after being transplanted and grown under SD conditions. FBD, represents the flower bud development stage, VC represents the visible color stage, EO represents the earlier opening stage, and OF represents the open-flower stage. C. Transcript level of *CmSVP* and *CmFTL3*, respectively, in 35S::*CmSVP* and VP64-*CmTPL1*-*2* transgenic chrysanthemums as well as wild type ‘Jinba’. Data represent the mean ± SEM of biological triplicates. Different letters represent a significant difference at P < 0.05 (one-way ANOVA with Fisher’s post hoc test).

#### CmSVP inhibits the transcription of *CmFTL3* by binding its promoter

To investigate whether CmSVP is capable of regulating the transcription of *CmFTL3* directly, we performed yeast one-hybrid (Y1H) assay. First, seven CArG elements of the *CmFTL3* genome sequence were selected as possible binding sites (P0-P6) (Fig. 2A). The results of Y1H showed that CmSVP could bind the full-length promoter of *CmFTL3* (Fig. 2B); then, the promoter was further segmented to confirm the interaction, which showed that CmSVP could interact with P1, P2, and P4 fragments, while P0, P3, P5, and P6 were the non-binding ones (Fig. S5). To further examine the CmSVP binding region of the *CmFTL3* genome sequence, ChIP-qPCR was used to screen the binding elements enriched by CmSVP. we found that CmSVP was able to target the CArG element in the promoter, 5’UTR, and intron regions. P0 and P3 are invalid sites for CmSVP binding, which is consistent with the results found in yeast (Fig. 2C, Fig. S5). As shown in Fig. S6, we detected the GFP-tag in the GFP fusion expression target protein CmSVP for ensuring the reliability of the assay. The EMSA assay with normal and mutation probes with the CArG motif in the promoter (P1 and mP1) and 5’UTR (P2, mP2; P3, and mP3) of *CmFTL3* suggested that CmSVP was also able to bind the *CmFTL3* promoter in vitro (Fig. 2D). The fragment of the sequence is shown in Fig. S7. The dual luciferase reporter assays by using 35S::*CmSVP* as an effector and *ProCmFTL3* as a reporter showed that the co-expression of the effector and reporter constructs reduced the *ProCmFTL3* activity effectively compared to the control groups (Fig. 2E). These results indicated that CmSVP could recognize and regulate *CmFTL3* directly in chrysanthemums.

**Fig. 2.**
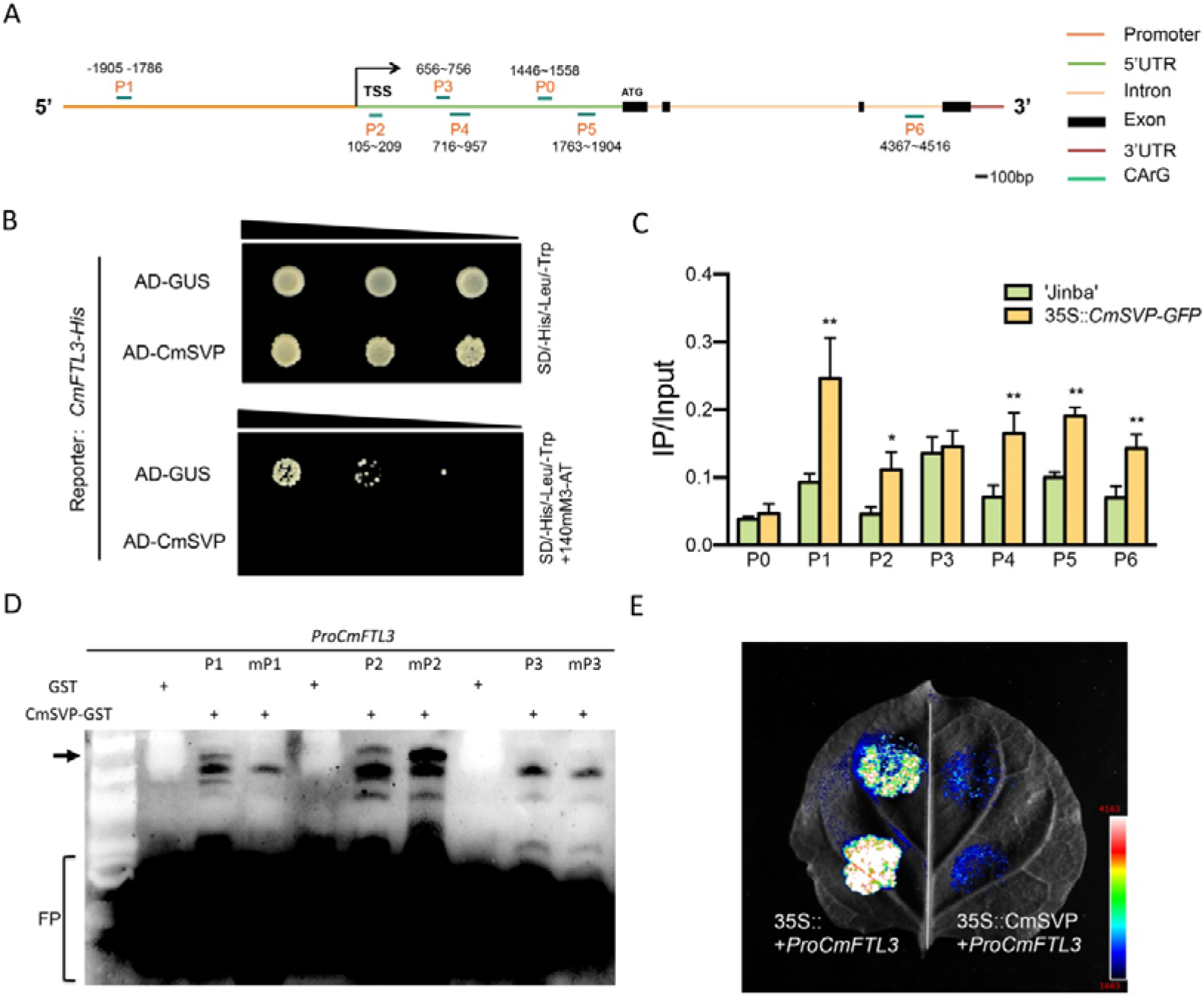
CmSVP directly binds to the CArG motif in the promoters of *CmFTL3*. A. Schematic diagrams showing the CmFTL3 genomic regions. The promoter is represented by an orange line, 5’UTR is represented by a green line, introns are represented by a light orange line, while exons are represented by black boxes. A flat ellipse (P0-P6) indicates the sites that have either single mismatch or are perfectly matched to the consensus binding sequence (CArG box) of MADS domain proteins. TSS, transcription start site. B. Interactions between CmSVP proteins and the promoters of *CmFTL3* in yeast cells. The 2249 bp fragment cloned in the promoter. pGADT7-GUS was used as a negative control. SD/-His/-Leu/-Trp indicates His, Leu, and Trp synthetic dropout media. 3-AT concentrations: 140 mM for *ProCmFTL3*. C. ChIP analysis of CmSVP binding to the regions of *CmFTL3* in the wild type ‘Jinba’ and transgenic lines of 35S::*CmSVP*-*GFP* chrysanthemums. Error bars indicate S.D. (n = 3 biological replicates). *P < 0.05 (Student’s *t*-test) for transgenic plants versus ‘Jinba’. D. EMSA of CmSVP binding to the P1/mP1, P2/mP2, and P3/mP3 fragment. ‘P1, P2, and P3 indicate labeled DNA probes, while mP1, mP2, and mP3 indicate mutated probes. Sequences are shown in Fig. S7. ‘+’ indicates presence and ‘-’ indicates absence. FP, free probe. E. Interactions of CmSVP proteins and the promoters of *CmFTL3* confirmed with dual luciferase reporter assays. The obtained sequence fragment of 2249 bp and P1 in *CmFTL3* promoter were detected as presented. 35S::+*ProCmFTL3* and 35S::+*proCmFTL3*-*P1* were used as controls.

### The interaction between CmSVP and CmTPL1-2

To elucidate the mechanism of CmSVP as a negative flowering regulator, a yeast two-hybrid (Y2H) assay was performed. With CmSVP as the bait protein, a cDNA fragment showing homology to the *Arabidopsis* TPL was identified, which was designated CmTPL1-2. Its characteristics were reported in our previous study(Zhang et al., 2019). CmSVP could interact with CmTPL1-2 in the yeast assay in vitro (Fig. 3A). CmSVP contains two EAR domains (LXLXLP), and we further confirmed that the interaction site of CmSVP-CmTPL1-2 occurred at the second EAR domain (LRLGLP) of the CmSVP (Fig. 3B). Moreover, firefly luciferase complementation imaging and co-immunoprecipitation (CO-IP) assay verified the interaction (Fig. 3C, D). Interestingly, the interaction between AtSVP and AtTPL in *Arabidopsis* was also confirmed in vitro and in vivo (Fig. S8), which indicated that the functions of SVP and TPL may be conserved in chrysanthemums and *Arabidopsis*.

**Fig. 3.**
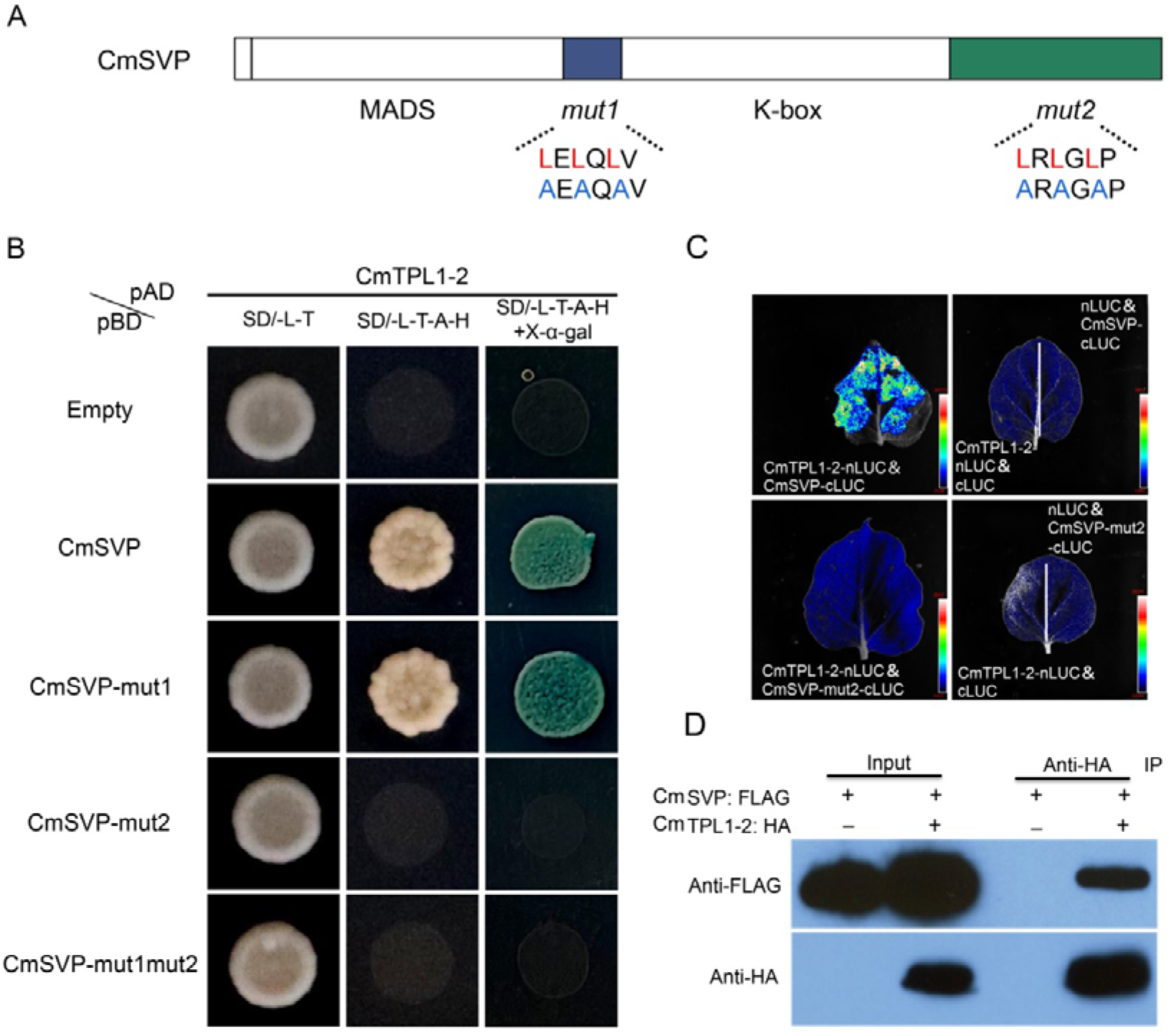
CmSVP interacts with CmTPL1-2. A. Schematic representation of the structure of CmSVP and highlighting the mutation of the two LXLXLX domains. B. The interactions were tested by yeast-two-hybrid. The yeast was transformed with empty vector or fusions of CmSVP, CmSVP-mut1, CmSVP-mut2, and CmSVP-mut1mut2 to the Gal4-binding domain (pBD), and fusions of CmTPL1-2 to the Gal4 activation domain (pAD). The yeast growth on nonselective (-L-T) and selective (-L-T-A-H without or with X-α-gal) SD medium. The second EAR motif LRLGLP was a determinant of the interaction between CmSVP and CmTPL1-2. C. FLuCI assay. Quantitative analysis of luminescence intensity showing the interaction between CmSVP and CmTPL1-2 in *N*. *benthamiana* epidermal cells. CmSVP/CmSVP-mut2 were fused to the C-terminal fragment of luciferase (cLUC), while CmTPL1-2 was fused to the N-terminal fragment of luciferase (nLUC). The interactions between nLUC and CmSVP/CmSVP-mut2-cLUC as well as CmTPL1-2-nLUC and cLUC were used as negative controls. Representative images of *N*. *benthamiana* leaves 72 h after infiltration are shown. D. Co-IP assay. CmTPL1-2-HA was pulled-down by immunoprecipitation of FLAG-tagged CmSVP. *N*. *benthamiana* leaves were agroinfiltrated with CmSVP-FLAG and CmTPL1-2-HA. Two days after agroinfiltration, total protein extracts were immunoprecipitated with an anti-FLAG antibody. CmTPL1-2-HA was detected in these fractions with an anti-HA antibody.

#### *CmTPL1*-*2* suppresses flowering time in chrysanthemums

In our previous study, the *TPL1-2* overexpression transgenics produced a higher number of rosette leaves and flowered around 15 days later than Col-0 (Zhang et al., 2019). The N176H mutation in the TPL of *Arabidopsis* is necessary and sufficient to induce the *tpl*-*1* mutant phenotype (Long et al., 2006; Szemenyei et al., 2008). We found that hsp::mut*CmTPL1*-*2* (N176H) lines would fail in floral repression when the 176^th^ amino acid of CmTPL1-2 was mutated from aspartic to histidine, which correlated with the late flowering phenotype compared to the Col-0 of *Arabidopsis* plants (Zhang et al., 2019). To further confirm the function of *CmTPL1*-*2* in chrysanthemums, each of three transgenic chrysanthemums, amiR-*CmTPL1*-*2*, and OX-*CmTPL1*-*2* (35S::*CmTPL1*-*2*) lines were selected after molecular identification (Fig. S8). Each of three independent overexpression lines (20-4, 22-3, and 25-1) flowered later than ‘Jinba’, while the amiR-*CmTPL1*-*2* (2-2, 4-2, and 9-7) and hsp::mut-*CmTPL1*-*2* (3-8, 4-2A, and 7-3) lines flowered significantly earlier than ‘Jinba’ (Fig. 4A, Fig. S9). At 15 days after transplantation and being grown under SD conditions, amiR-*CmTPL1*-*2* plants were already exhibiting differentiation of the involucral primordial, which took place 12–15 days earlier than the WT plants. Moreover, the amiR-*CmTPL1*-*2* plants that had already reached the VC represented the visible color stage, while the OX plants were still 10–13 days away from the flower buds emerging phase, which was 22–25 days later than that in the WT plants (Fig. 4B). We then investigated whether the expression of *CmFTL3* was changed in *CmTPL1*-*2* transgenic chrysanthemums. We found that the levels of *CmFTL3* mRNA in 35S::*CmTPL1*-*2* plants showed a dramatic decrease, but exhibited an increase in amiR-*CmTPL1*-*2* plants compared with the WT ‘Jinba’ (Fig. 4C). These results confirmed the interaction between CmTPL1-2 and CmSVP genetically.

**Fig. 4.**
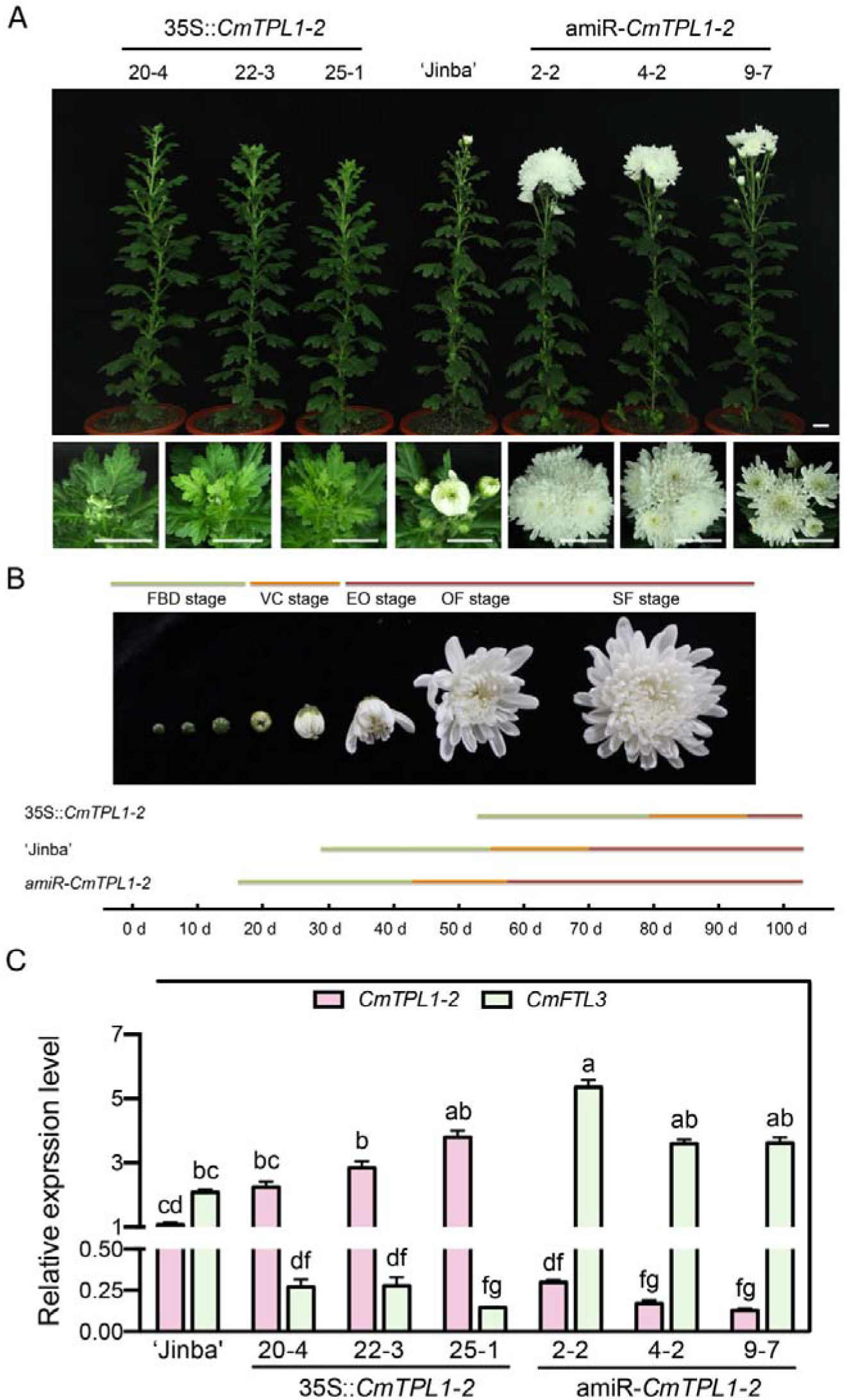
Phenotypes of *CmTPL1*-*2* transgenic ‘Jinba’ plants. A. The phenotypic consequence of overexpression (20-4, 22-3, and 25-1) and knocking down (2-2, 4-3, and 9-7) of *CmTPL1*-*2*. Scale bar = 1.5 cm. B. Developmental process of flower buds in wild type ‘Jinba’ and transgenic plants. Flowering time was calculated after being transplanted and grown under SD conditions. FBD represents the flower bud development stage, VC represents the visible color stage, EO represents the earlier opening stage, and OF represents the open-flower stage. C. The transcript level of *CmTPL1*-*2* and *CmFTL3*, respectively, in 35S::*CmTPL1*-*2* and amiR-*CmTPL1*-*2* transgenic chrysanthemums. Data represent the mean ± SEM of biological triplicates. Different letters represent a significant difference at P < 0.05 (one-way ANOVA with Fisher’s post hoc test).

#### CmSVP recruits CmTPL/CmTPR to repress *CmFTL3* in the floral transition of chrysanthemums

Because the TPL functions as a co-repressor, it is suggested that the transcription repression level regulates *CmFTL3* by *CmSVP* requiring *CmTPL1*-*2*. To investigate this, a reporter and effector vector construction was used for a transient assay in chrysanthemum protoplasts. The sequence of 5’UTR and the promoter of *CmFTL3* was fused with the *LUC* reporter gene, respectively (Fig. 5A). The 5’UTR-*CmFTL3* and *proCmFTL3* were strongly repressed by CmSVP in ‘Jinba’, and CmSVP presented stronger repression activity in the *CmFTL3* promoter compared with 5’UTR. Therefore, the protoplast isolated from amiR-*CmTPL1*-*2* transgenic chrysanthemum was used for transcription activity detection. Moreover, the activity was partly repressed, which may be due to the incomplete disappearance of CmTPL1-2 and the existing homologous genes of CmTPL/CmTPR (Fig. 5B). Therefore, the obtained mutation of *CmTPL1*-*2* (N176H) refers to mut*CmTPL1*-*2*, which was present in loss-of-function of all CmTPL/CmTPR family members, and was used for transcription activity detection. The results showed that 5’UTR and the promoter of *CmFTL3* remained rarely changed in mut*CmTPL1*-*2* with a heat shock vector (pMDC30) transgenic 3-8 line compared to the WT ‘Jinba’ after 37°C treatment (Fig. S10). The results suggested that the CmSVP-dependent repression of *CmFTL3* in the loss of CmTPL/CmTPR function in chrysanthemums was impaired.

**Fig. 5.**
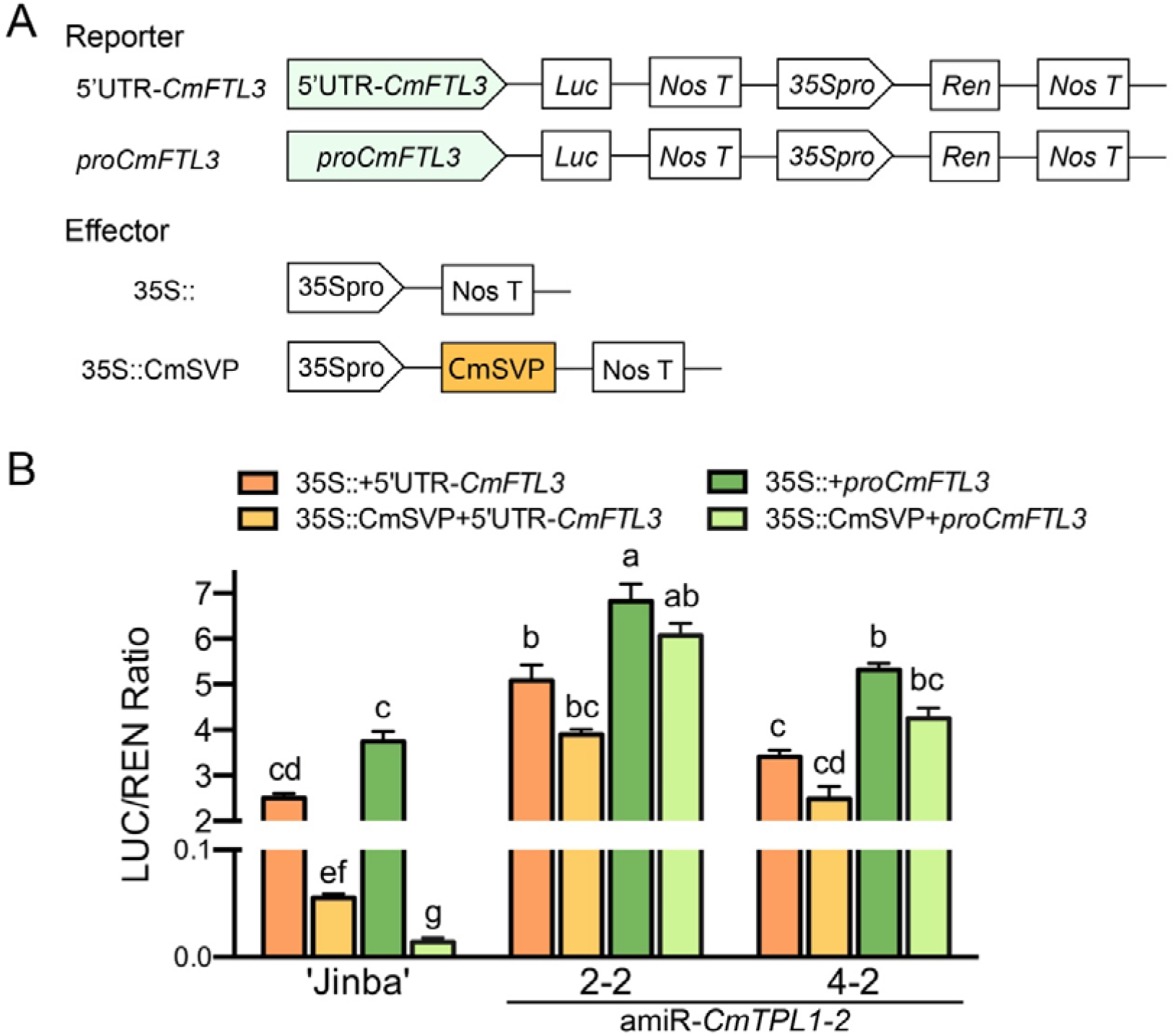
CmSVP suppresses *CmFTL3* transcription mediated by CmTPL/CmTPR activity. A. The construction diagram of the reporter 5’UTR-*CmFTL3* and *proCmFTL3* and effectors. B. Transient expression analysis in protoplasts of chrysanthemum genetic transformation lines (amiR-*CmTPL1*-*2*). Repression of *CmFTL3* by CmSVP was mostly dependent on CmTPL1-2. The LUC/REN is the average ratio of the bioluminescence of firefly luciferase to that of firefly luciferase. Data represent the mean ± SEM of biological triplicates. Different letters represent a significant difference at P < 0.05 (one-way ANOVA with Fisher’s post hoc test).

#### Dependence of *SVP* relies on *TPL*/*TPR* in the regulation of flowering is conserved in chrysanthemum and *Arabidopsis*

To determine the conserved function of SVP recruiting for TPL to suppress its target gene expression, we further performed the related assay in *Arabidopsis* mutant *tpl*-*1*, which acts as a type of dominant negative allele for multiple TPL/TPR family members. 35S::*AtSVP tpl*-*1* and 35S::*CmSVP tpl*-*1* were generated by introducing the *AtSVP* and *CmSVP* gene driven by the 35S promoter, respectively, with molecular identification (Fig. S13). Neither *AtSVP* nor *CmSVP* could revert the phenotype of early flowering that was caused by *tpl*-*1*, but both *AtSVP* and *CmSVP* overexpression in the WT plants Col-0 delayed flowering significantly (Fig. 6).

**Fig. 6.**
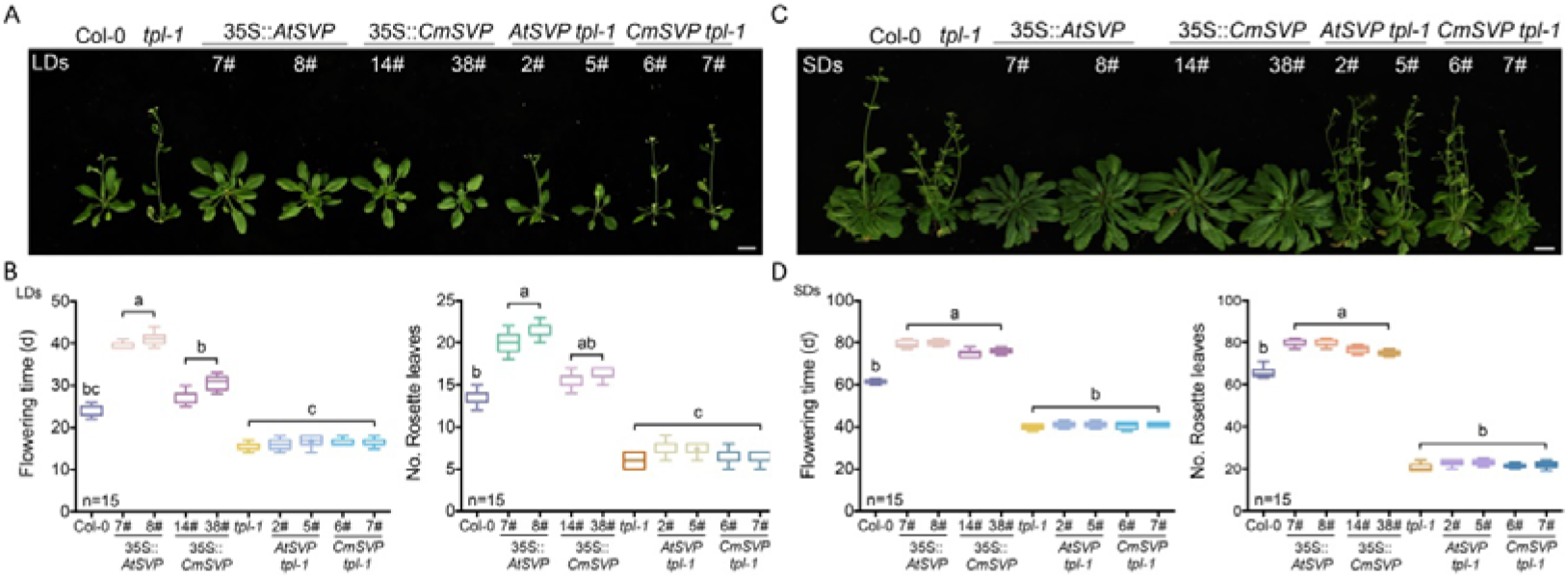
Flowering characterization of *AtSVP* and *CmSVP* overexpression plants, respectively, in the Col-0 and *tpl*-*1* background. A. Phenotype of wild type Col-0, *tpl*-*1*, and transgenic lines in LDs. Scale bar = 1.5 cm. B. Statistics of wild type Col-0, *tpl*-*1*, and transgenic lines in LDs. C. Phenotypes of wild type Col-0, *tpl*-*1*, and transgenic lines in SDs. Scale bar = 1.5 cm. D. Statistics of wild type Col-0, *tpl*-*1*, and transgenic lines in SDs. Data represent the mean ± SEM of biological triplicates. Different letters represent a significant difference at P < 0.05 (one-way ANOVA with Fisher’s post hoc test). Meanwhile, the late flowering effect of 35S::*CmSVP* and the early flowering effect of 35S::*CmSVP*-*VP64* in *Arabidopsis* reveal that *CmSVP* is a flowering inhibitor; in addition, *CmSVP* was able to fully rescue the *svp*-*31* mutant of Col-0 (Fig. S11), which genetically suggests that the regulatory function of *CmSVP* in the flowering time of *Arabidopsis* and chrysanthemum is conservative. *FT* is expressed in companion cells of leaf phloem tissues(Chen et al., 2018). To specifically reveal the function of TPL, the construct SUC2::*mAtTPL/*SUC2::*mCmTPL1*-*2* (AtTPL/CmTPL1-2 N176H), which expressed the *tpl*-*1* mutant protein driven by a *SUCROSE-PROTON SYMPORTER 2* (*SUC2*) phloem companion cell-specific promoter from *Arabidopsis*, was transformed into *Arabidopsis* Col-0 and mutant *svp*-*31*, respectively, with molecular identification (Fig. S13). We found that the three lines of SUC2::*mAtTPL svp31* (10#, 4#, and 11#) and SUC2::*mCmTPL svp31* (4#, 1#, and 38#) had a similar earlier flowering phenotype with *svp*-*31* plants either in LDs or SDs compared with the Col-0 WT plants (Fig. 7), which was consistent with that of the double mutant *tpl*-*1 svp*-*31*, which had similar early flowering compared with the single mutant (Fig. S12). The molecular identification showed in Fig. S14. These results confirmed that the interaction between TPL and SVP is genetically conserved in *Arabidopsis* and chrysanthemums, and that *SVP* requires TPL/TPR to function as a floral repressor.

**Fig. 7.**
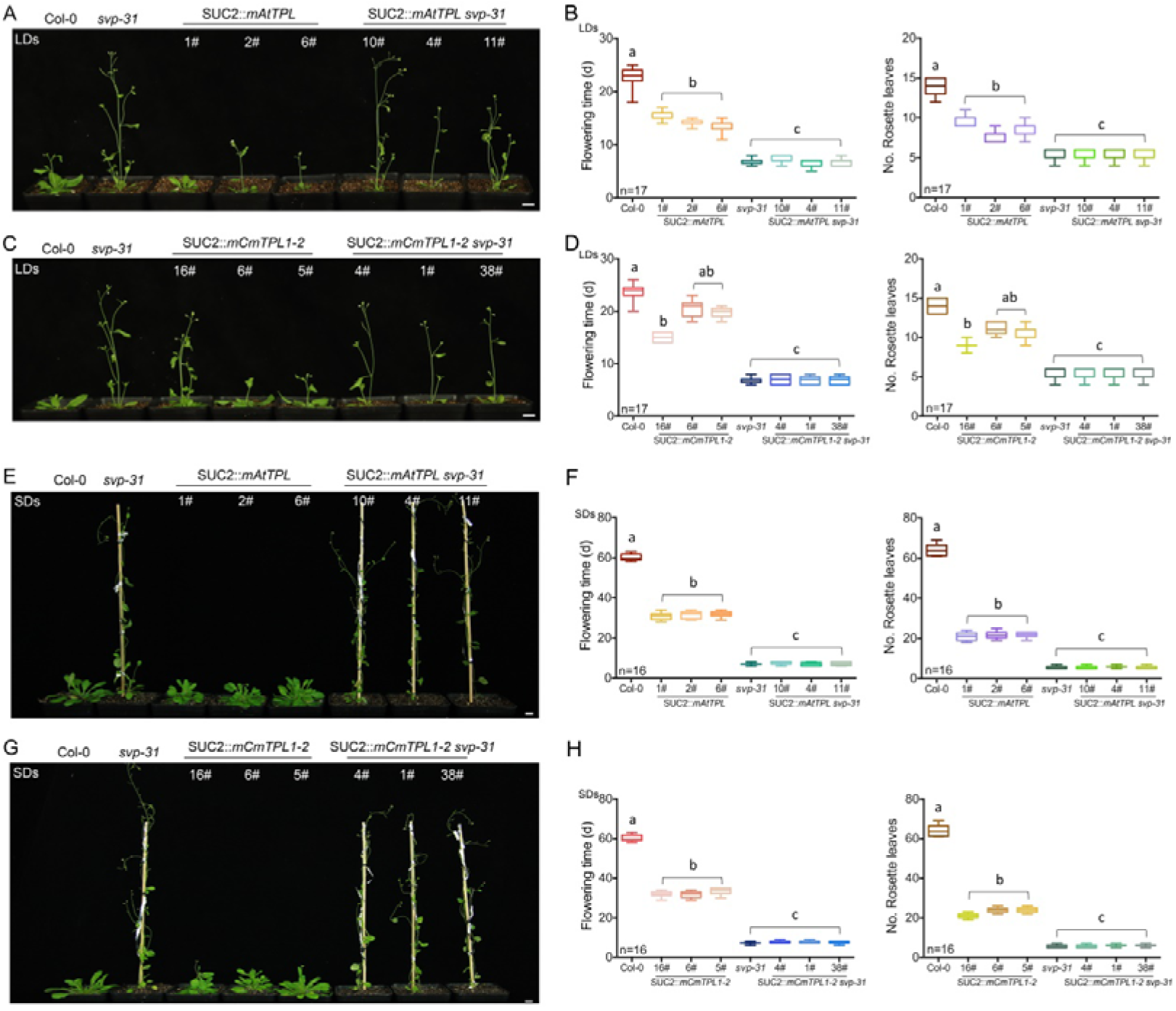
Flowering time of SUC::*mAtTPL* and SUC::*mCmTPL1*-*2* transgenic plants in Col-0 and *svp-31* backgrounds on long and short days. A and C. Representative images of SUC::*mAtTPL/CmTPL1*-*2*/Col-0 and SUC:: *mAtTPL/CmTPL1*-*2*/*svp*-*31* plants and their parental genetic background (Col-0 and *svp*-*31*) under LDs at flowering. Scale bar = 1.5 cm. B and D. Quantification of flowering time (B) and rosette leaf number (D) during long day photoperiods. E and G. Representative images of SUC::*mAtTPL/CmTPL1*-*2*/Col-0 and SUC:: *mAtTPL/CmTPL1*-*2*/*svp*-*31* plants and their parental genetic background (Col-0 and *svp*-*31*) under SDs at flowering. Scale bar = 1.5 cm. F and H. Quantification of flowering time (F) and rosette leaf number (H) during short day photoperiods. Data represent the mean ± SEM of biological triplicates. Different letters represent a significant difference at P < 0.05 (one-way ANOVA with Fisher’s post hoc test).

## Discussion

In this study, the mechanism of SVP in repressing the floral transition in both *Arabidopsis* and chrysanthemums through the interaction with a co-repressor protein TPL was elucidated. There was a conserved EAR motif in SVP proteins which was required for the interaction (Fig. 3), SUC2::*tpl*-*1* transgenic *Arabidopsis*, or amiR-*CmTPL1*-*2* transgenic chrysanthemum, attenuates the function of SVP as a transcriptional repressor (Fig. 5,6,7). SVP mediates flowering responses through many pathways, which respond by perceiving signals from different endogenous and environmental factors, such as the GA and thermosensory factors (Gregis et al., 2013; Lee et al., 2013). Recent studies on *Arabidopsis* have demonstrated that GA promotes the expression of *FT* and *SOC1* by suppressing a group of flowering repressors (*ELF3*, *SVP*, *TEM1*, and *TEM2*) via the *GAF1-TPR* complex (Fukazawa et al., 2021). The loss of SVP in *Arabidopsis* suggested that *SVP*-mediated control of the expression of *FT* evolved in the course of the transition from the vegetative to reproductive phase, thus, tracking the changes at the ambient temperature (Lee et al., 2008; Lee et al., 2013; Song et al., 2013). As small non-coding RNAs (microRNAs) act as significant regulatory role during flowering of plants. Studies revealed that miR172 is regulated by SVP and SPL9 (SQUAMOSA PROMOTER BINDING PROTEIN-LIKE 9) of *Arabidopsis* reproduction period (Lee et al., 2010; Zhen et al., 2012). With the gradual in-depth study of SVP homologous genes in plants, the structure and function of SVP have become clearer, but the mechanism of its inhibitory function has not yet been revealed (Gregis et al., 2010; Mauren et al., 2014). Here, the results showed that a SVP-TPL transcriptional complex suppressed *FT* to limit the floral transition during the vegetative phase (Fig. 8).

**Fig. 8.**
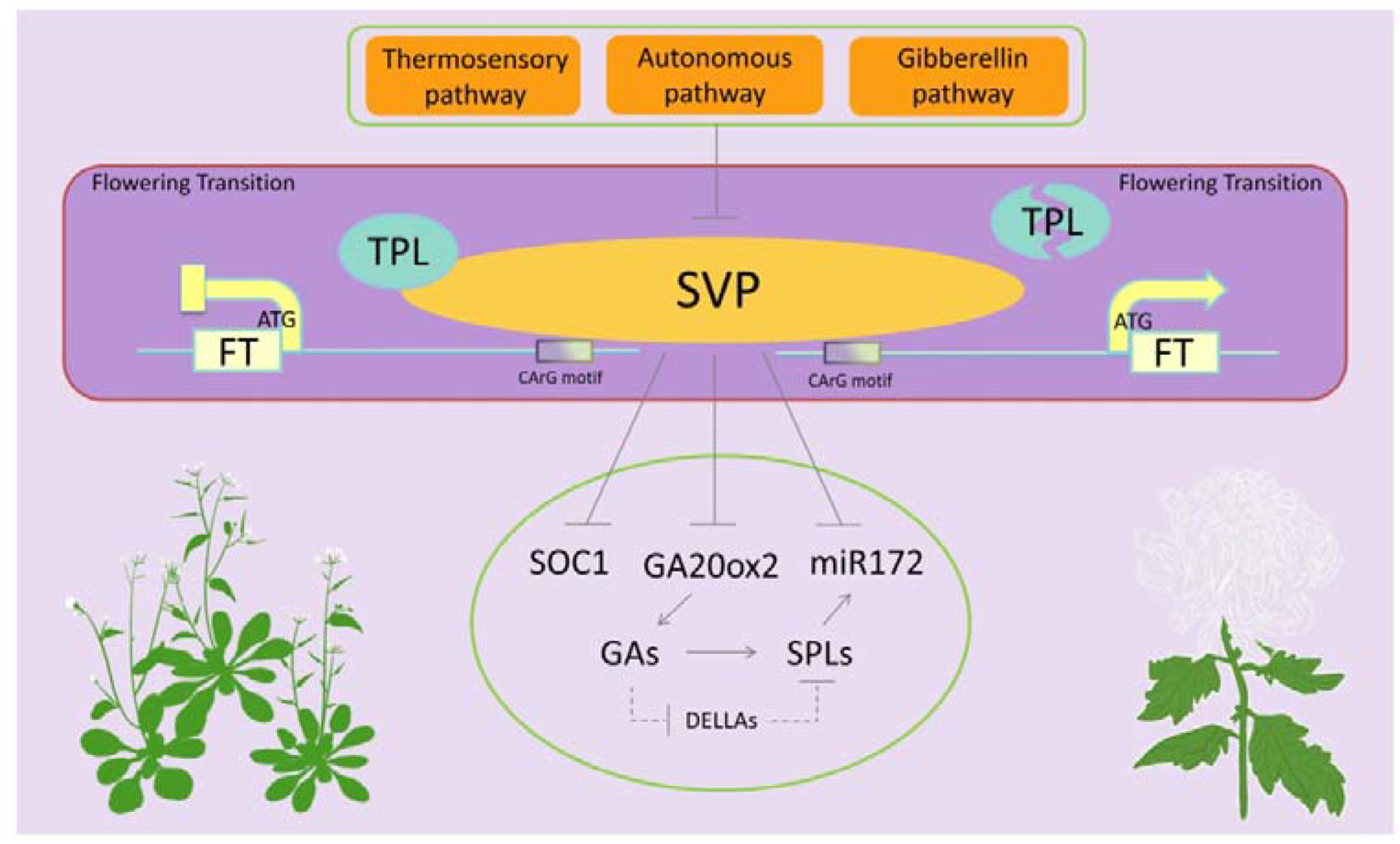
Schematic representation of SVP and TPL mediating the integration of flowering signals. *SVP* expression is inhibited by the thermosensory, autonomous, and gibberellin (GA) pathways in *Arabidopsis*. Downregulation of *SVP* transcription contributes to increased the expression of *SOC1*, *GA20ox2*, and miR172. Higher GA levels increases the SPLs transcription and release SPLs proteins by DELLAs repression. Arrows and bars indicate promoting and repression effects, respectively. The interactions proposed in this study was shown in the dark purple box with the red outline, SVP transcription factor represses a key flowering regulator *FT* by binding to the CArG motif of its promoter both in *Arabidopsis* and chrysanthemum. However, SVP need recruit TPL to complete the process of inhibition. In the absence of TPL, the action path of SVP-FT for flowering will be ineffective.

SVP has a typical plant-specific restriction EAR domain (LXLXLX); however, whether the motif is required for its transcriptional repression activity is unclear (Li et al., 2008; Lisha et al., 2011). For *AtSVP* and *CmSVP*, which were isolated from *Arabidopsis* and chrysanthemums, respectively, the second EAR domain (LRLGLP) of the sequence is essential for interacting with TPL. When it is mutated, the action relationship is invalid and irrelevant to the first EAR domain (Fig. 3). As the conserved EAR motif inhibits transcription processes, the specific function of the first EAR (LELQLV) is unknown.

The SVP transcription factor binds to the promoters of *FT*, which completely reverse the effect in triggering an early flowering response(Li et al., 2008). We verified that the process is highly conservative in chrysanthemums (Fig. 2). The two most studied *Arabidopsis* transcriptional repressors for flowering are FLOWERING LOCUS C (FLC) and SVP, which are both present in MADS-box protein(Andrés et al., 2014; Song et al., 2013). FLC can form dimers and function with other redundant MADS-box proteins to suppress flowering by repressing the transcriptions of floral activators, such as *FT* and *SOC1* (Searle et al., 2006). In *Arabidopsis*, the CArG motif (284-302 bp) in the first intron of *FT* can be strongly enriched by FLC proteins, but SVP’s binding appears to be weaker than that of FLC (Helliwell et al., 2006; Lee et al., 2007). Studies on SVP have revealed that it is able to bind with the CArG motif (−1235-1225 bp) directly in the *FT* promoter, and this is also the only site on the promoter that exercises the inhibitory function(Lee et al., 2007; Song et al., 2013). This is somewhat different from the site of action in chrysanthemums. Here, we showed, CmSVP was able to target the CArG element in the promoter, 5’UTR, and intron regions, as well as in the promoter. This indicated that the regulation through binding to CArG was conserved.

Transcriptional co-repressors play considerable roles in entrenching the adequate levels of gene expression during flowering (Plant et al., 2021). As conserved co-repressors, TPL/TPR was filtered as a partner of SVP in this study. Some previous studies have shown that the floral transition in precise regulation requires TPL/TPR running at multiple points in the pathway to flowering (Espinosa-Ruiz et al., 2017; Fukazawa et al., 2021; Plant et al., 2021; Tao and Estelle, 2018). And moreover, most studies of TPL have been carried out on model plant *Arabidopsis* but not on non-model plants. Here, overexpression of *AtSVP* and *CmSVP* in the *TPL*/*TPR* loss-of-function mutant *tpl*-*1* showed futility in delaying flowering (Fig. 6). Moreover, the recruiting dependency relationship between SVP and TPL either in *Arabidopsis* or chrysanthemum was conserved. Experiments in chrysanthemum protoplasts prove the dependence of CmSVP on CmTPL/CmTPR in the process of inhibiting *CmFTL3* transcription (Fig. 5). SVP acts as a core flowering repressor that always functions along with other potent transcription factors (Golembeski and Imaizumi, 2015; Zhen et al., 2012). In *Arabidopsis*, J3 with a typical modular sequence of the J-domain, which encodes a DnaJ-like heat shock protein and often appears as a protein chaperone, represses SVP activity to induce *SOC1* and *FT* expression (Shen et al., 2011; Shen and Yu, 2011). CO interacts with microproteins miP1a/b, TPL, and JMJ14 to prevent flowering in the shoot apical meristem (SAM) until the leaf-derived FT protein triggers the transition to the reproductive growth phase. However, the study did not detect the previously identified TPL/TPR-interacting repression domain containing transcription factors and the formation of a higher order repressor complex is a small process that might be subject to the surrounding conditions (Rodrigues et al., 2021). SVP can interact with TERMINAL FLOWER 2/LIKE HETEROCHROMATIN PROTEIN 1 (TFL2/LHP1) to regulate floral patterning(Liu et al., 2009), while TFL2 recognizes H3K27me3 to repress the expression of many genes including *FT* (Liu et al., 2018). TPL typically associates with histone deacetylase (HDAC) in planta (Krogan et al., 2012), and also interacts with histone-binding protein MSI4, CHROMATIN REMODELLING 4 (Larsson et al., 1998; Turck et al., 2007). This reflects the organizational complexity of flowering transition in plants.

As the flowering process in plants is an extremely complex and delicate event and partly accounts for the occurrence of different species and evolutionary adaptation, the crosstalk of SVP with other factors that can ensure the reproductive success remains unknown. Therefore, the interaction between SVP and TPL revealed in this study may represent a rising molecular interconnection among the respective families of conserved regulators, which is linked intermediately to flowering.

## Materials and Methods

### Plant materials and growing conditions

The experiments were centered on the *C*. *morifolium* cultivar (cv.) ‘Jinba’, which was obtained from the Chrysanthemum Germplasm Resource Preserving Centre (Nanjing Agricultural University, China). Vegetatively propagated cuttings at the 5–6 leaf stage were grown in a 1:1 mixture of garden soil and vermiculite under a 16 h photoperiod (day/night temperature regime of 23°C/18°C, relative humidity 70%). The *Arabidopsis* plant stocks employed were WT ecotype Col-0 which from The Arabidopsis Information Resource (www.arabidopsis.org/). And the mutants *svp*-*31* (*SALK_026551*) and *tpl*-*1* (*At1g15750*) provided respectively by Xu’ lab and He’s lab. All *Arabidopsis* plants were soil-grown under a constant temperature of 22 ± 2°C, a 16 h photoperiod, and 70% relative humidity.

### Isolation and analysis of the gene sequence

Total RNA was extracted using the RNAiso reagent (TaKaRa, Tokyo, Japan) from snap-frozen chrysanthemum leaves of ‘Jinba’, as recommended by the manufacturer. A 1 μg aliquot of RNA was used for the synthesis of the cDNA first strand using a PrimeScriptTMRT reagent Kit containing gDNA eraser (TaKaRa, Shiga, Japan). The cDNA was used as the template and primers in TableS1 were used to PCR amplify the sequence. The amplicon was inserted into the pMD-19T vector (TaKaRa, Tokyo, Japan) by T4 DNA ligase (TaKaRa, Tokyo, Japan) for sequencing. Mutant primers (Table S1) of *CmTPL1-*2 were designed based on the site of the 531 bases (A) to be mutated to C on the website (http://www.bioinformatics.org/primerx/cgi-bin/DNA_3.cgi), which has been described previously (Zhang et al., 2019). The *CmSVP* site was mutated from (TTATTTAAGAAAGCTGAAGAG) to (TTTAAAAAGGCCGAGGAG), which is able to bind miR396(Yang et al., 2015).

### Transactivation activity assay

LR Clonase™ II enzyme mix (Invitrogen, Carlsbad, CA, USA) was used to recombine with pGBKT7 (Clontech, Mountain View, CA, USA) and pGBKT7-VP16 vector (provided by Teng’s lab) for the construct. Studies have shown that fusion with VP16 (a fragment of viral DNA sequence encoding the peptide DALDDFDLDML) is capable of turning a repressive transcription regulator into an activator (Guo et al., 2018; Suzuki et al., 2014; Triezenberg et al., 1988). A transactivation activity assay was performed following the manufacturer’s instructions. Salmon sperm DNA carrying pGBKT7-*CmSVP* (BD-CmSVP), pGBKT7-VP16-CmSVP (VP16-CmSVP), pGBKT7 (BD, negative control), GAL4 and pGBKT7-VP16 (BD-VP16, positive control) inserted into the yeast strain Y2H (Clontech, Mountain View, CA, USA), which next transferred to the SD/-Trp medium. After 3 days, a single clone was selected for cultivating and transferred to the SD/-Trp-His medium with 0 or 20 mg/mL X-α-gal. According to the directions, if the protein possesses transcriptional activation activity, it should bind to the GAL4-BD upstream promoter sequence of *His3* as the GAL4-BD could regulate the *His3* expression. As a result, yeast colonies grew on SD/-Trp-His and turned blue on SD/-Trp-His with X-α-gal.

### Subcellular localization

The full-length coding region (minus the termination codon) was amplified with appropriate modifications, generating 35S::GFP-CmSVP. It was then transformed into protoplasts of wild-type ‘Jinba’ after incubation for 12–14 h at 28°C. GFP fluorescence was then detected using a confocal laser scanning microscope (LSM800, Zeiss).

### Yeast two-hybrid assay

The coding regions of *CmTPL1*-*2* was amplified and cloned into pGADT7, and *CmSVP* or *CmSVP*-mut1/mut2 was amplified and cloned into pGBKT7 (Clontech) for Y2H assays. The Yeastmaker Yeast Transformation System 2 was used according to the manufacturer’s instructions (Clontech). pGBK-53 and pGADT were used as positive controls, while pGBK-Lam and pGADT were used as negative controls. All combinations were transferred to the SD/-Leu-Trp medium by the yeast strain Y2H (Clontech, Mountain View, CA, USA), and then transferred to the SD/-Leu-Trp medium. Yeast colonies were grown on SD/-Leu-Trp-His-Ade and turned blue on SD/-Leu-Trp-His-Ade containing X-α-gal if an interaction existed between the proteins.

### RNA isolation and gene expression analysis

Samples were taken when the transgenic material and WT were grown till 18–20 leaves appeared under LD conditions (16 h/8 h photoperiod) and then transferred to SD conditions (8 h/16 h photoperiod) for 3 days. Meanwhile, SAM and mature leaves were selected, and samples were taken in the early morning (*FT* gene expression was the highest). Quantitative primer of *CmFTL3* is designed according to the sequence provided by http://172.30.0.105/VIROBLAST/VIROBLAST. PHP sequence design The cDNA was used as the template for qRT-PCRs based on Fast SYBR Green Master Mix (www.bimake.com). The qRT-PCR involved an initial denaturation (95°C/2 min), followed by 40 cycles of 95°C/15 s, 60°C/15 s, and 72°C/15 s. The reference sequence was the *Actin8* gene for *Arabidopsis* and *C*. *nankingense EF1α* gene for chrysanthemums, and the relative transcript abundances were calculated using the 2^−ΔΔCT^ method (Livak and Schmittgen, 2001). The set of qRT-PCR primer sequences used is listed in Table S3.

### *Arabidopsis* transformation

The p35S::*CmSVP*, p35S::*AtSVP*, 35S::*CmSVP*-*VP64*, p35S::*CmTPL1*-*2*, p35S::*AtTPL1*-*2*, SUC2::*mCmTPL1*-*2* and SUC2::*mAtTPL1-2* transgenes were introduced into *Arabidopsis* by the *A. tumefaciens* strain EHA105. 1/2 of the MS medium, which contained 50 μg mL^−1^ hygromycin or 1 μg mL^−1^ kanamycin, was applied for transformed progeny selection. Each of three independent T3 transgenic plants was obtained and validated by using the PCR primer pair in Table S1 and S2 for amplification.

### Co-immunoprecipitation assay

Co-IP assays were conducted at He’s Laboratory (Li et al., 2018). Briefly, the total protein was extracted from tobacco expressing CmTPL1-2-HA and CmSVP-FLAG using anti-HA affinity gel (E6779, Sigma), followed by western blotting with anti-FlAG (A8592, Sigma), and anti-HA (12013819001, Roche).

### Yeast one-hybrid assay

The CDSs of CmSVP were inserted into the pGADT7 vector to generate the recombined construct pGADT7-CmSVP, while the CDS of GUS (β-glucuronidase) was inserted into the pGADT7 vector as the negative control. The *CmFTL3* promoter and 5’UTR fragments were cloned into the pHIS2 vector. The primer pairs used for gene cloning are listed in Supplementary Table S1. Subsequently, all constructs were transformed into Saccharomyces cerevisiae strain Y187 using the lithium acetate method. Subsequently, yeast cells were inoculated on a selective medium lacking Trp, Leu, and His (SD/-Trp/-Leu/-His). The selected colonies were then inoculated on a -Trp/-Leu/-His medium supplemented with an appropriate concentration of 3-AT and grown for 3 days at 28°C, the binding was identified by spot assay.

### Dual-luciferase reporter assay (leaves of *N*. *benthamiana*)

The fragment of the *CmFTL3* promoter (http://172.30.0.105/viroblast/viroblast.php) was cloned into the pGreenII0800-Luc vector, which contained a reporter gene encoding firefly luciferase (kindly provided by Dr. Huazhong Shi, Texas Tech University, Lubbock, TX). *A. tumefaciens* strain GV3101 harboring *CmFTL3*::LUC, p35S::GFP-*CmSVP*, p35S::GFP-*CmTPL1*-*2*, and p35S::GFP was grown in infiltration medium (2 mM Na_3_PO_4_, 50 mM MES, 100 mM acetosyringone) to an OD600 of 0.5 and then introduced via a syringe into the leaf of a 4–5-week-old *Nicotiana benthamiana* plant. After 48–96 h, a CCD camera was used to observe luciferase activity.

### Dual-luciferase reporter assay (protoplasts of chrysanthemums)

Overall, 10 μg plasmid of *proCmFTL3*-*P1*, 35S::*CmSVP*-Flag, 35S::*CmTPL1*-*2*-Flag, and 35S::Flag were transformed to amiR-*CmTPL1*-*2* and 35S::*CmTPL1*-*2* (heat shock induced vector, pMDC30) transgenic lines of chrysanthemum protoplasts. After 40% PEG-mediated transformation, the protoplasts were placed in a dark environment at 24°C for 20 h. It should be emphasized that 35S::*CmTPL1*-*2* (heat shock induced vector, pMDC30) can be expressed only after treatment with 37°C, and wild-type ‘Jinba’ of chrysanthemum was used as the control. The Renilla and firefly luciferase activities were measured using a Dual-luciferase Reporter Assay System (Promega, cat. # e1910).

### Electrophoretic mobility shift assay (EMSA)

The fusion proteins of CmSVP were generated through prokaryotic expression in vitro. The CDSs of CmSVP were cloned into the PGEX-5T vector containing a Gst (GST) target to generate recombined vectors. Then, these recombined vectors were transformed into *Escherichia coli* BL21 (DE3). IPTG was used to induce protein production. The fusion proteins were purified using the MagneGST™ Pull-Down System (Promega). The subsequent EMSAs were performed using a LightShiftTM Chemiluminescent EMSA Kit (Thermo Fisher, New York), following the manufacturer’s instructions. The GST protein was used as a negative control, and unlabeled probes were used for probe competition. The resulting samples were loaded onto a pre-run native 6.5% polyacrylamide gel using TBE buffer as the electrolyte. After electro-blotting onto a nylon membrane (Millipore, Darmstadt, Germany) and UV cross-linking (2000 J for 5 min), the membrane was incubated in blocking buffer for 30 min and rinsed in washing buffer. Finally, a CCD camera was used to visualize the chemiluminescent signal.

### ChIP-qPCR assay

The transgenic chrysanthemum p35S::GFP-*CmSVP* was applied to the ChIP-qPCR assays. Moreover, the EpiTect ChIP OneDay Kit (Qiagen) was used according to the manufacturer’s instructions. A GFP-specific antibody was used in the assay. Subsequent quantitative real-time PCR (qRT-PCR) used sequence-specific primers, which are provided in Table S3.

## Acknowlegements

This work was financially supported grants from the National Natural Science Foundation of China (31872146), and a project funded by the Priority Academic Program Development of Jiangsu Higher Education Institutions. We thank Dr. Yuehua Ma (Central laboratory of College of Horticulture, Nanjing Agricultural University) for assistance in using ultra high resolution confocal microscope (LSM800, Zeiss, Germany).

## Author contributions

J.F.J., F.D.C, and Z.X.Z. conceived and designed the experiments; Z.X.Z., H.Q. and Y. Q.Z. performed most of the experiments; G.Z., E.L.S, G.F.L, W.X.L and X.R.C. provided technical support; Y.H.H, S.B., S.M.C, W.M.F. and Z.Y.G. provided conceptual advice; G.Z., H.Q. and Z.X.Z. contributed to the Co-IP assay; E.L.S. and H.Q. contributed to the ChIP assay; H.Q., Z.X.Z., Y.Q.Z. and R.Q.H contributed to plants transformation; Z.X.Z., H.Q. and J.F.J. analyzed the data and wrote the manuscript.

